# Harnessing the power of poplar tree natural genetic variation for the development of future sustainable biofuels and bioproducts: a droughted marginal-land experiment for multi-disciplinary investigations

**DOI:** 10.1101/2024.01.11.575272

**Authors:** Gail Taylor, Jack H Bailey-Bale, Marie C Klein, Suzanne Milner, Jin-Gui Chen, Wellington Muchero, Peter Freer-Smith, Timothy J. Tschaplinski, Jerry Tuskan

## Abstract

The emerging bioeconomy offers significant potential to replace fossil-fuel-based energy, manufacturing, and processing with that utilizing biomass as the raw feedstock. However, feedstock production from non-food crops such as fast-growing trees, must be delivered at scale, in a reliable and consistent manner, utilizing marginal land unsuitable for food crops and with minimum inputs. This new generation of feedstock crops has a limited history of domestication. Foundational knowledge is required to enable rapid selection and breeding for improved cultivars and varieties to enable large-scale planting of 600M ha, globally over the coming decades. Here, we describe an innovative field platform with over 1,000 unique genotypes of fast-growing poplar (*Populus trichocarpa*) trees, each sequenced and being subjected to a controlled drought. The 6.5 ha site provides opportunities to bring together multi-disciplinary phenotyping science linked to computational, and AI approaches, enabling the link between complex plant traits and their underlying genes to be rapidly established and translated into the development of improved climate-resilient germplasm for a future at-scale bioeconomy.

## Introduction

Biomass feedstock from fast-growing woody plants will play an important role in the emerging bioeconomy, defined as the use of biological resources to generate goods, services, and energy. Many nations are developing bioeconomy strategies as a crucial part of a more sustainable future, including the USA (The Whitehouse, 2023). Lignocellulosic crops have the potential to deliver sustainable feedstock for multiple uses, including liquid aviation fuel, heat, and power generation, linked to carbon capture and storage, as well as the supply of multiple high-quality chemicals and replacements for petroleum-based products (IEA Bioenergy, 2022), enabling reductions in greenhouse gas emissions.

Relative to the use of fossil fuels, which still heavily dominate the global energy sector, bioenergy crops can aid decarbonization efforts. It has been estimated that global resources for biomass to supply these future demands are significant and could require up to 600M ha of land (Intergovernmental Panel on Climate Change, 2022); however, many of these model estimates are focused on the deployment of Bioenergy with Carbon Capture and Storage (BECCS), and thus the requirement for land and supply of feedstocks for other non-food uses associated with a fully integrated biorefinery is likely to be significantly higher. For a successful transition to a sustainable circular bioeconomy, operating within the ecological boundaries of the planet, there is a significant challenge for this emerging and paradigm-changing industry to supply substantial amounts of feedstock that is both environmentally and economically sustainable, and deliverable in an equitable way to local and global communities. Adverse effects on land competition for food should be avoided, with limited negative impacts on biodiversity and water availability. To achieve this, new lignocellulosic biomass crops are required to be able to grow with minimal inputs, particularly water, on marginal land (Intergovernmental Panel on Climate Change, 2022).

One of the fastest-growing woody plants on the planet and a source of biomass feedstock for the future is *Populus* - poplar and aspen, which are found over much of the northern hemisphere (Taylor, 2002). Their potential was recognized when the North American species *Populus trichocarpa* (black cottonwood), was the first tree species to be sequenced (Tuskan, et al., 2006), shortly after the model herbaceous plant Arabidopsis (The Arabidopsis Genome Initiative, 2000), providing an initial glimpse of the genetic resource that could be harnessed for the future. The tumbling costs of DNA sequencing, improved technologies, and increased depth of genome coverage and future technological advances (Shendure et al., 2019), mean that plant traits can now more easily be resolved to the level of the gene. Alongside enhanced computational power, the potential of AI and the development of new innovative approaches in a number of contrasting technologies, from gene editing (Zhang et al., 2018) to cheap sensors in the environment (Levintal et al., 2022), improved satellite imagery, linked to plant function (Tattaris et al., 2016) and high throughput phenotyping (Tardieu et al., 2017), these new technologies are converging to drive a step change in the rate of novel discoveries on the foundational links between plant performance and genomic control. This is of critical importance to the emerging bioeconomy, and of particular interest is the scale-up and production of resilient biomass resources that can grow in resourcelimited environments with few water and nutrient inputs; land undesirable for food production, and delivering few ecosystem services (Khanna et al., 2021). These marginal landscapes demand new insight into the link between environment, plant performance, and genes, and here we describe a new resource that addresses this problem.

Of the three major bioenergy feedstock tree species, poplar (*Populus* sp.), willow (*Salix* sp), and eucalypts (*Eucalyptus* sp.), the knowledge and technology foundation for genetic improvement is most advanced in poplar, with *Populus* sp. widely dispersed across the northern hemisphere, reflecting its origin in Eurasia and multiple colonizations across other regions including North America (Liu et al., 2022). Of the thirty or so poplar species, vastly different environments have been colonized, including extremes of Northern Sweden, characterized by low temperatures and short days for *P. tremula*, to the African sub-continent and dry arid zones for *P. euphratica*.

This breadth and depth of genetic diversity across poplar species globally is a treasure trove that can be harnessed for the future. A rich genetic resource such as this strengthens the likelihood of a species developing tolerance to multiple biotic and abiotic stresses, highlighting the importance of quantifying, harnessing, and conserving adaptive traits for future selection, breeding programs, and biotechnological design to enable climate-smart bioenergy forests fit for multiple purposes. This is likely to include feedstocks for Sustainable Aviation Fuel (SAF), in addition to energy with Carbon Capture and Storage (BECCS), alongside nature-based solutions to sequester carbon below ground. Each of these pathways requires a deeper biological understanding than that currently available. For drought tolerance in particular, broad diversity exists and has been quantified in physiological, biochemical, and morphological characteristics in poplar that have facilitated its adaptation to droughted natural environments (Monclus et al., 2006; Street et al., 2006), and thus it seems likely that these traits are highly tractable and can be elucidated for the future development of improved feedstocks.

Poplar seeds require nearly immediate germination; therefore seedbanks are a challenging method for maintaining and preserving germplasm. This said, they are however easily propagated from hardwood cuttings, which has enabled the establishment of several common garden trials globally, where germplasm is clonal and an immortal population can thus be established, and this can have major benefits for maintenance, scale-up and supply of plant material, which will be required in the coming decades. This propensity for easy regeneration of clonal material has been utilized in a limited way to explore natural genetic diversity across several *Populus* species, detailed in Table 1.

**Table 1:** Natural populations of *Populus*, collected and planted in common garden experiments and used to elucidate the links between phenotype and genotype, providing foundational knowledge to underpin future selection, breeding, and biotechnology routes to improvement. Modified from Allwright (2017).

| Species | Location | Notes |
| --- | --- | --- |
| *P. nigra* | UK / Belgium | 948 genotypes drawn from multiple riparian populations across western Europe (France, Spain, Germany, Italy, Netherlands, and Hungary) and planted in a common garden in Northington, England (51.1°N) and Savigliano, Italy (44.6°N), used to unravel genetic basis of bioenergy traits (Allwright et al., 2016). |
| *P. nigra* | USA | Population includes 599 genotypes drawn from 17 different open-pollinated families collected from Italy, Belgium, and Serbia and established at a test plot in Boardman, Oregon (45.8°N) in 2008. Genotyped using 433 SNPs in 39 candidate genes for chemical wood properties by (Guerra et al. 2013). Rapid *within-gene* linkage disequilibrium (LD) was observed, suggesting a close association between marker SNPs and causative polymorphisms. |
| *P. nigra* | China | 134 genotypes sampled from across 15 European countries (Romania, Yugoslavia, Spain, Italy, Bulgaria, Croatia, Turkey, Russia, UK, Netherlands, Belgium, Czech Republic, Slovakia, Germany, and Hungary) and established at 3 field sites in Beijing, inner Mongolia, and Shaanxi (Ding & Nilsson, 2016). |
| *P. trichocarpa* | USA | 1100 genotypes established at multiple field sites in the USA. 1052 genotypes from *P. trichocarpa*'s core range in northern Oregon, Washington, and southern British Columbia, with an additional 48 from northern California, central Oregon, and central British Columbia. Utilized by Slavov et al., (2012) in their study of the geographical structure and LD in *P. trichocarpa* and subsequently by Evans et al (2014) to determine the genomic signature of adaptive trait associations (Evans et al., 2014). |
| *P. tremula* | Sweden | The Swedish Aspen (SwAsp) Collection – 116 genotypes taken from 12 populations across Sweden from 56°- 66°N and planted in 2 common gardens in Ekebo and Savar at 55.9° and 63.4°N, respectively (Escamez et al., 2023; Robinson et al., 2012), the latter to study bioenergy feedstock traits |

**Table 2:**
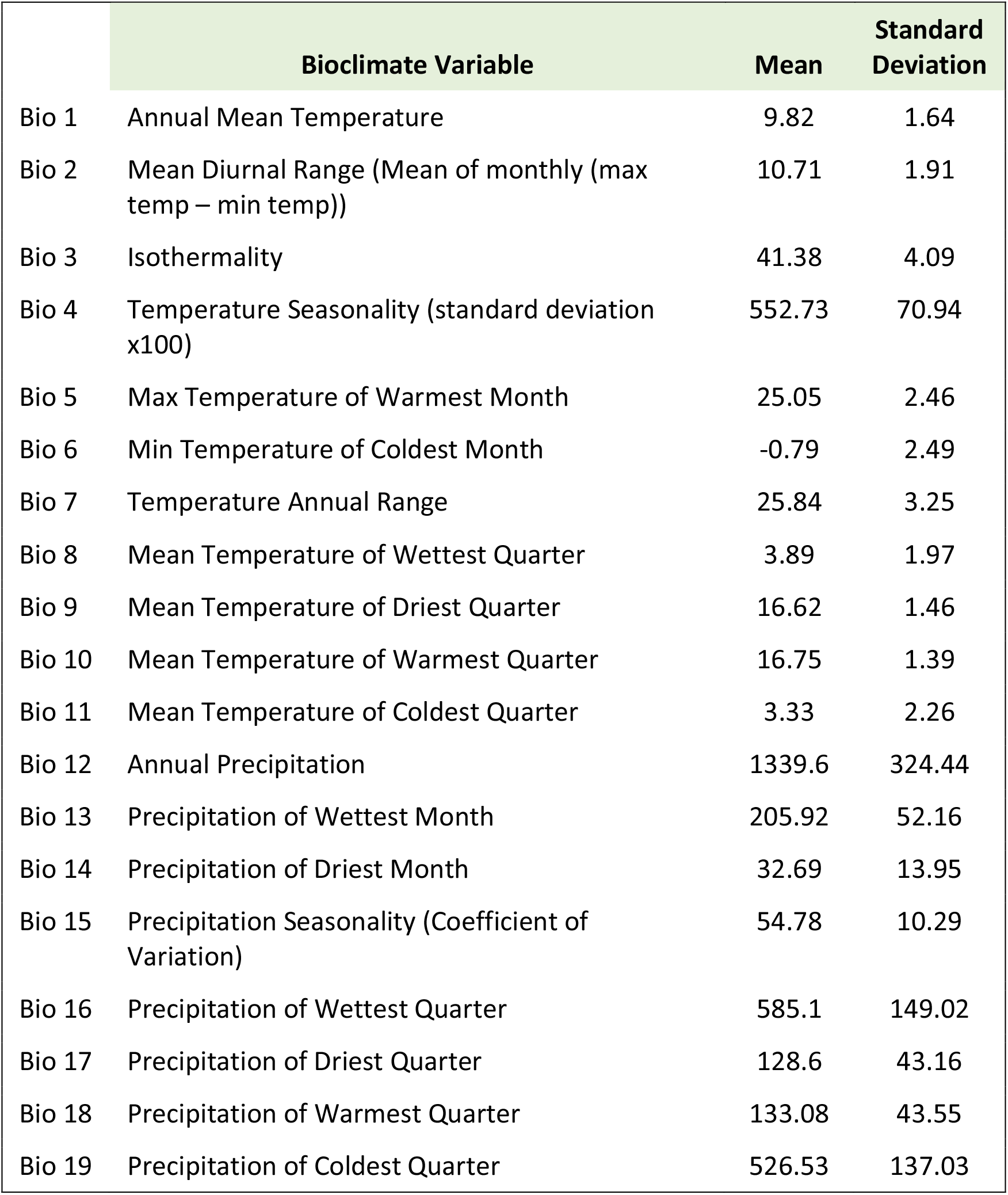
Bioclimatic variables (mean and standard deviation) are shown among *P. trichocarpa* genotypes describing climate of origin. Data derived from Worldclim (Fick and Hijmans 2017).

|  | Bioclimate Variable | Mean | Standard Deviation |
| --- | --- | --- | --- |
| Bio 1 | Annual Mean Temperature | 9.82 | 1.64 |
| Bio 2 | Mean Diurnal Range (Mean of monthly (max temp – min temp)) | 10.71 | 1.91 |
| Bio 3 | Isothermality | 41.38 | 4.09 |
| Bio 4 | Temperature Seasonality (standard deviation x100) | 552.73 | 70.94 |
| Bio 5 | Max Temperature of Warmest Month | 25.05 | 2.46 |
| Bio 6 | Min Temperature of Coldest Month | -0.79 | 2.49 |
| Bio 7 | Temperature Annual Range | 25.84 | 3.25 |
| Bio 8 | Mean Temperature of Wettest Quarter | 3.89 | 1.97 |
| Bio 9 | Mean Temperature of Driest Quarter | 16.62 | 1.46 |
| Bio 10 | Mean Temperature of Warmest Quarter | 16.75 | 1.39 |
| Bio 11 | Mean Temperature of Coldest Quarter | 3.33 | 2.26 |
| Bio 12 | Annual Precipitation | 1339.6 | 324.44 |
| Bio 13 | Precipitation of Wettest Month | 205.92 | 52.16 |
| Bio 14 | Precipitation of Driest Month | 32.69 | 13.95 |
| Bio 15 | Precipitation Seasonality (Coefficient of Variation) | 54.78 | 10.29 |
| Bio 16 | Precipitation of Wettest Quarter | 585.1 | 149.02 |
| Bio 17 | Precipitation of Driest Quarter | 128.6 | 43.16 |
| Bio 18 | Precipitation of Warmest Quarter | 133.08 | 43.55 |
| Bio 19 | Precipitation of Coldest Quarter | 526.53 | 137.03 |

Growing poplar from distinct geographic locations of origin in shared environment field trials has enabled the effects of genetic diversity to be isolated and compared between populations. This has facilitated the study of specific traits of interest, for example, pest resistance, yield, and seasonality, to name a few, and some work on *P. nigra* has been undertaken with respect to drought tolerance (Allwright, 2017; Taylor et al., 2021). The experimental power of these trials is now, however, greatly enhanced with recent technology developments, as highlighted above. Utilizing these modern techniques across hundreds of genotypes, this paper details the methodology we are employing in order to elucidate the genetic basis of drought tolerance and yield in *Populus trichocarpa*.

## Materials and Methods

The Center for Bioenergy Innovation (CBI), is one of four U.S. Department of Energy Bioenergy Research Centers (https://genomicscience.energy.gov/bioenergy-research-centers/), funded to accelerate the development of sustainable biofuels and bioproducts. At the University of California, Davis, a 6.5 ha field site was established in 2020, focusing on the rapid domestication of the bioenergy-relevant, woody perennial tree species, *Populus trichocarpa* (*P. trichocarpa*), where we aim to harness natural genetic variation to create superior feedstock plants. To achieve this, in January 2020 ∼17000 hardwood *P. trichocarpa* cuttings from wild trees, originating from across their natural range (Figure 2, with variable rainfall and temperature shown) were received by UC Davis from Oak Ridge National Laboratory. These dormant cuttings, consisting of 1382 unique genotypes, were collected from trees grown in a *P. trichocarpa* common garden in Corvallis (OR). The hardwood cuttings were briefly stored in a 5 °C cold room before being transferred to the greenhouse. Each cutting was dipped into a solution of water and Hormodin 2 before being inserted into 30 cm leach tubes, each filled with Premier Pro HP Mycorise 3.8 CF. All benches we soaked daily with fertilizer water for 58 days; more than 90 % of the hardwood cuttings were successfully established. Following this, the trees were transferred for 14 days in a lath house located within 1 km of the experimental site prior to being planted. A 2.5 ha drought site and a 4 ha control site were prepared, each containing 3 replicate blocks. Under the challenging conditions of the COVID-19 pandemic, we devised a safe protocol for genotype planting. From 1382 *P. trichocarpa* genotypes, we selected lines with at least 6 healthy saplings, each of which was assigned a block – control block 1, 2, 3 or drought block 1, 2, 3. On April 10^th,^ 2020, including guard trees around the perimeter of all 6 experimental blocks, over 7000 poplar saplings were planted at 8 ft x 10 ft, spacing. Establishing and then maintaining the health of the trees and site has been an important factor in the success of this unique genetic resource. For each tree, a 1 m2 black ground mat was used to reduce competition from weeds and each tree was also placed into a 50 cm high translucent protective tube. These were effective at deterring small mammals present at the field site from eating young buds and foliage from the saplings. Fully controlled irrigation was deployed with each row of trees, supplied by 16 mm drip line irrigation, with an emitter positioned every 30 cm. Throughout the 6 experimental blocks, the associated irrigation risers were equipped with K-Rain BL-KR V2.0 controllers. 8 controllers operate across the 6 experimental blocks, enabling fine manual operation of the water supplied to trees. The controllers enabled the design and adjustment of irrigation programs based on the day, water budget, start time, and start duration. The hourly flow rate of irrigation from each riser on the site was also measured to enable fine control of irrigation. Soil moisture status was monitored at the site using both Instrotek 503 ELITE Hydroprobe and WATERMARK soil moisture sensors with associated monitors, enabling close monitoring of both gravimetric and volumetric soil moisture content. Hydroprobe measurements were made every 10-14 days in each block at 3 depths; 30 cm, 60 cm, and 120 cm. Two access tubes, one inserted in the planting row and the other between the planting rows, allowed us to sample directly underneath the irrigation lines and at the furthest point from them. Volumetric water content was collected hourly by 7 tensiometer sensors in each treatment, collecting and storing the data locally for monthly download.

During the establishment year in 2020, all experimental blocks were fully irrigated enabling successful site establishment. Due to the extremely arid weather conditions experienced in Davis, this meant a whole-site adjustment of the watering schedules to accommodate the long, hot, and dry summer days. From April 2021, the drought blocks were subjected to an increasing and monitored soil moisture deficit, by restricting irrigation across the 3 experimental drought blocks relative to that applied across the control blocks, with effective results (Figure 2), where soil water potential in the drought treatment, was held between 0.1-0.15 MPA in both 2021 and 2022 (Figure 2), providing an extreme long-term contrast to the fully irrigated treatment. Careful monitoring of the soil moisture and tree health was important to ensure the fine balance of mildly water-stressing the trees was achieved, whilst simultaneously avoiding mass dieback throughout the drought blocks. Constant updates to the irrigation schedule were important, considering local weather conditions and the progression of the tree’s evaporative demand as they accumulated foliage. The same strategy was employed in 2022, the second year of water stress deployment. Between October 2021 and March 2022 field site irrigation was switched off as winter rains recharged the groundwater. We monitored tree phenology to identify indicators that the trees were breaking dormancy, and in April 2022, we deployed the second year of drought to the three drought blocks.

## Results and Discussion

The CBI *Populus trichocarpa* population at UC Davis comprises a diverse range of genotypes originating from diverse climates, which exhibit substantial genetic diversity across their range (Figure 2, Table 1). The distribution of these genotypes across the Northwest Pacific region provides an opportunity to study genetic and physiological mechanisms underlying adaptation to varying climatic conditions, including differences in temperature, precipitation, latitude, and elevation. A total of 1382 trees were examined in the study (Figure 2). A principal component analysis of bioclimatic variables (Fick & Hijmans, 2017) reflecting the location of origin for the trees, reveals that PC1 (which accounts for 45% variance), distinguishes cool-wet from warm-dry environments. Thus, we expect that adaptation along this gradient will reflect divergence in tolerance to water limitation (Blumstein et al., 2020).

The field site is located in Davis, CA, which experiences a Mediterranean-type climate, characterized by distinct and contrasting seasons, with hot, dry summers and cool, wetter winters, making it a valuable and reliable setting for studying drought responses, as water limitation can be achieved simply by restricting irrigation during summer months which experience little to no rainfall. In contrast, a control “well-watered” treatment was achieved by continued irrigation, serving as a reference for comparison with the drought-treated trees.

The objective of this experiment is to determine the physiological and morphological responses of the trees to drought stress and to identify the potential mechanisms that trees use to cope with water scarcity. To ensure that the water-limitation strategy here was effective, soil moisture deficit was measured across the blocks. As expected, the drought treatment saw a consistently decreasing soil water potential over the summer, while the control treatment maintained a high soil water potential (Figure 2).

The reduction of growth performance, apparent by eye in the field, is validated in estimates of biomass (height, tree diameter, etc) (Figure 2). Reduced soil moisture is associated with a decrease in biomass, which suggests drought tends to have a significant impact on productivity, inspiring future analyses of genotype-by-environment interactions to identify genotype susceptibility to drought (ΔBiomass = Biomass_Drought_ - Biomass_Control_). Similar analyses with ecophysiological traits, such as phenology, stomatal density, and water use efficiency, will provide further details of genetic diversity for plasticity in response to drought and the potential contributions of specific traits to drought tolerance.

The extensive phenotyping conducted over multiple seasons at both control and drought sites is yielding valuable data on the physical and biochemical traits associated with drought tolerance in *P. trichocarpa* (Figure 1). Moreover, the genetic variability of this population has been characterized by whole genome resequencing (McKown et al., 2014; Tuskan, Difazio, et al., 2006). Combined with phenotypic data, genome-wide association studies (GWAS) are underway to identify genetic variants that contribute to traits and performance under drought and control treatments (Figure 3).

**Figure 1:**
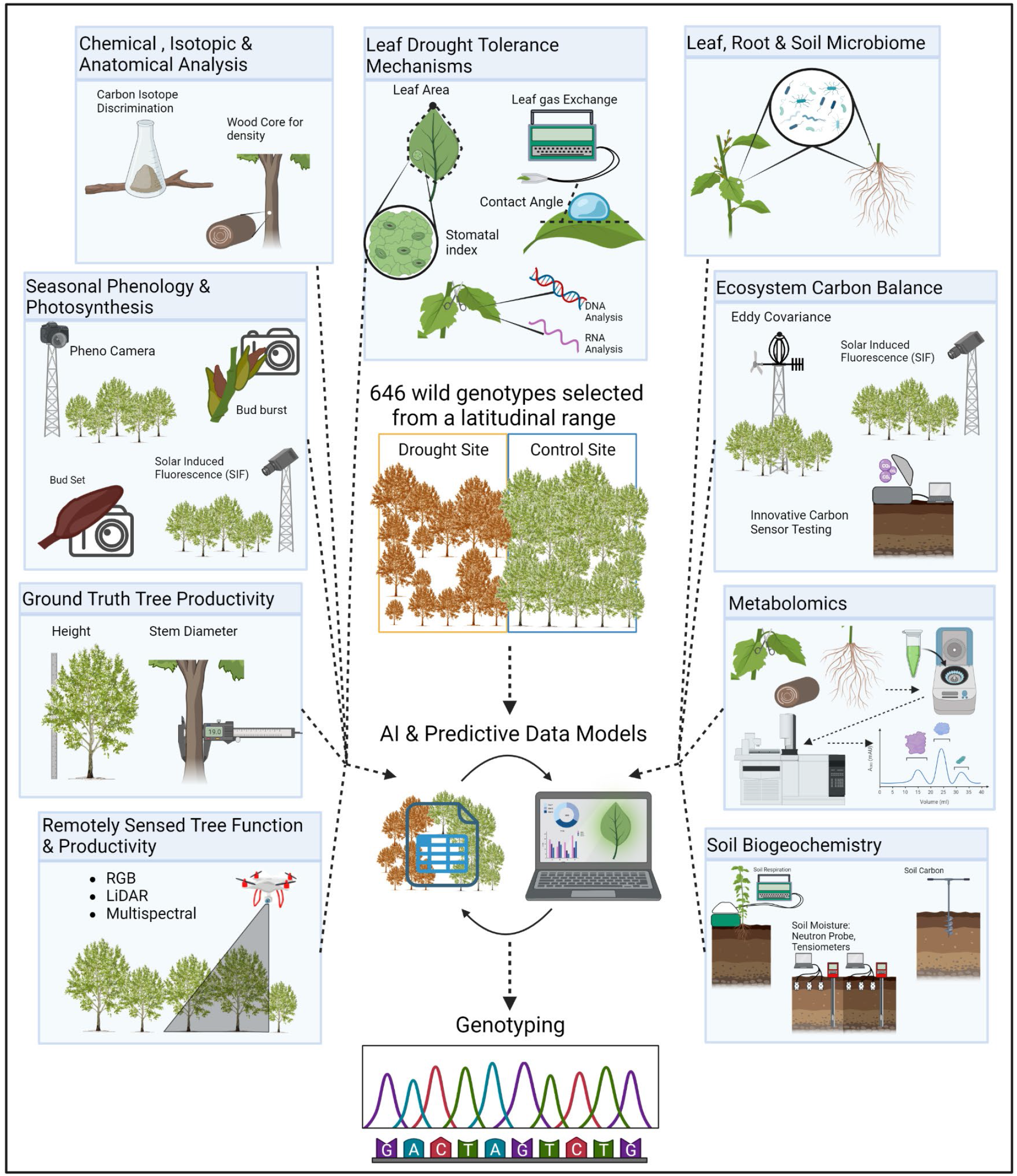
Harnessing the power of natural genetic variation. Over 1,000 unique genotypes, were replicated in each of 6 experimental blocks, selected from a latitudinal gradient from the western USA, with contrasting rainfall, and planted in a fully replicated experiment with controlled irrigation enabling defined ‘drought’ and ‘control’ treatments. At approximately 6.5 ha, the site is a platform enabling multi-disciplinary research groups to test a plethora of hypotheses on the genetic basis of poplar drought tolerance at scales ranging from metabolic and cellular to whole canopy and ecosystem function. This enables data-driven predictions on gene models linked to traits of interest that can be further explored through gene editing and genomic selection to design trees for droughted, marginal land.

**Figure 2.**
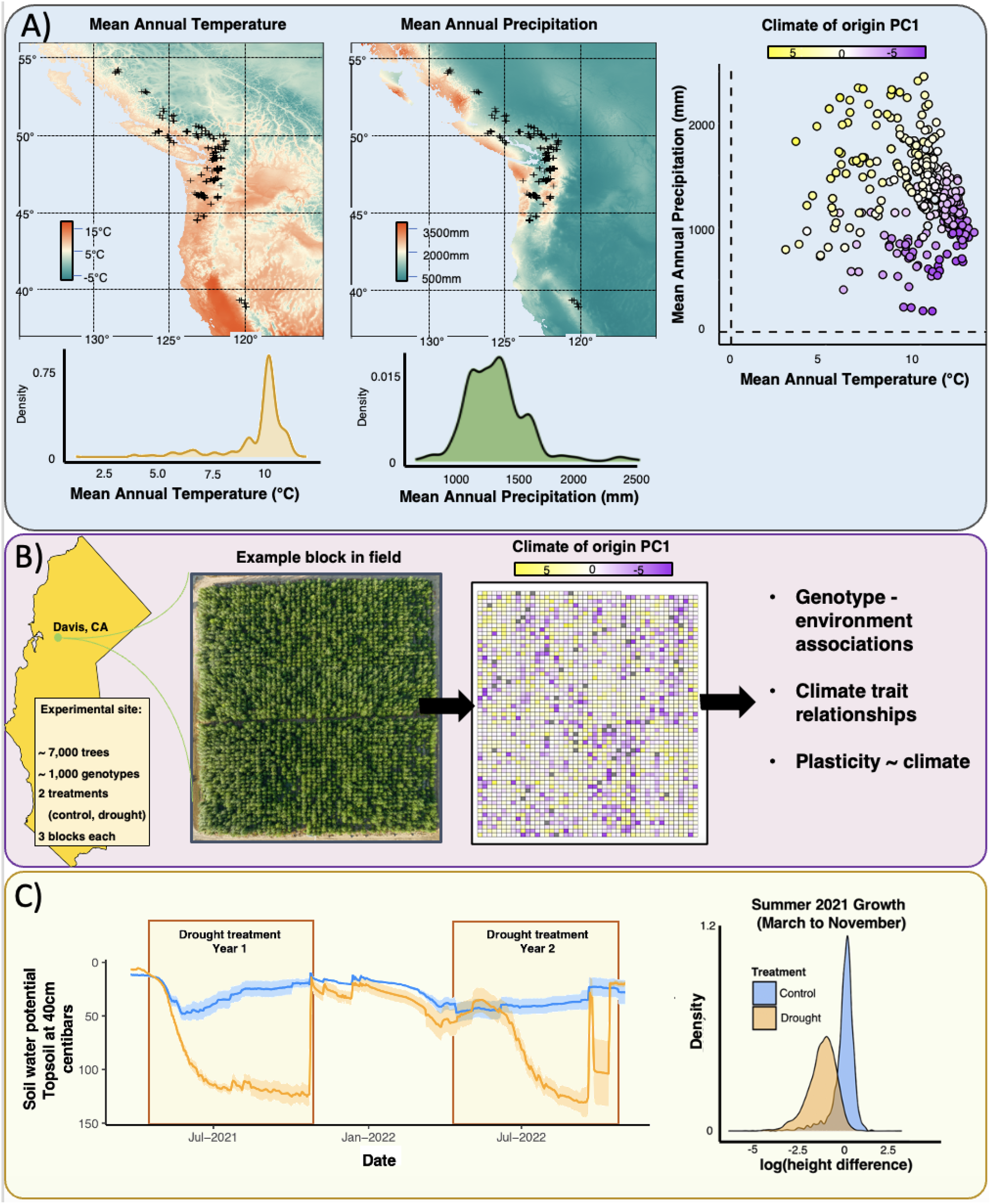
Overview of the experiment. **A)** Geographical range and bioclimatic variation of *Populus trichocarpa* along the Pacific Northwest, with climates summarized into principal components. B) Field trial to study variation in response to drought by diverse genotypes. An example of block 1 with genotypes randomized is shown. C) Soil moisture measurements show the drying down and rehydration of the drought blocks relative to the control in 2021 and 2022. This resulted in phenotypic responses, such as height differentiation between the treatments.

**Figure 3.**
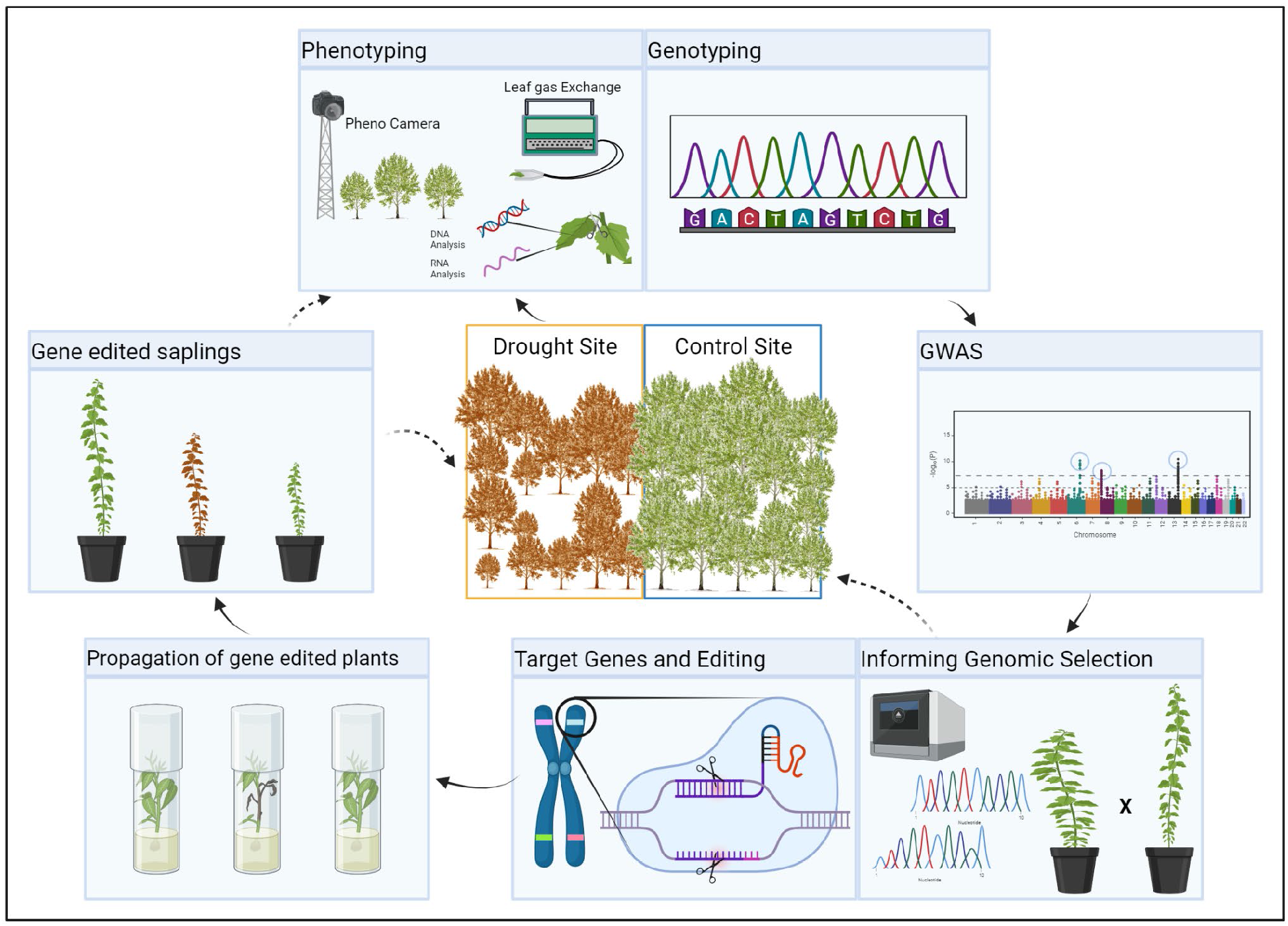
Phenotyping and genotyping data from the site trees is enabling genomic targets that underpin traits of interest to be rapidly identified and tested in the gene editing pipeline, followed by further ‘proof of concept’ testing of modified trees back in the field.

Genetic variants identified through these experiments may yield candidates for genome-enabled breeding, such as marker-assisted selection, genomic selection strategies, and genome editing. Quantifying the effect sizes on the multi-environment performance of loci at genomic scales can better inform genomic selection and the selection of markers that are associated with desirable traits to accelerate the breeding process. Moreover, variants with strong associations to traits and a clear functional effect of the candidate genes can inform applications of gene editing technologies like CRISPR/Cas9, to rapidly deploy desirable drought tolerance alleles in trees with other agronomically desirable traits (Ahmed et al., 2021; Maharajan et al., 2022; Nascimento et al., 2023).

This study provides a novel system for measuring potentially adaptive traits and collecting empirical data to test evolutionary and ecological models of adaptation. A better understanding of the genetics and traits responsible for adaptation to warmer-drier environments may improve the prediction of eco-evolutionary responses to climate change, with implications for conservation and ecosystem functioning (e.g., carbon cycling) in the Anthropocene (Exposito-Alonso et al., 2018; Monroe et al., 2018).

In summary, the CBI UC Davis poplar field platform is an essential tool for advancing our understanding of plant biology and enabling the supply of improved *P. trichocarpa* germplasm as a source of fast-growing trees suited to marginal land for the emerging bioeconomy.

## Conclusions

We describe a new manipulative, large-scale common garden experiment where approximately 1000 unique genotypes of the biomass feedstock tree, *P. trichocarpa*, selected from areas of contrasting rainfall are being grown, subjected to drought, and intensively observed. This site on marginal land with limited rainfall and significant heat is enabling research to unravel the genetic basis of biomass supply for the future bioeconomy and the potential of poplar to deliver in such extreme environments. The platform provides an opportunity for multi-disciplinary research to ensure a systems-level approach, providing insights into the genetic basis of traits from molecular and cellular, through to ecosystem functioning. Such interdisciplinary collaboration, combined with innovative approaches, including genomic editing, will enable the delivery of bespoke biomass as a valuable resource for the emerging bioeconomy.

## Acknowledgements

We thank the UC Davis Department of Plant Sciences Field crew, and multiple undergraduate student interns. Funding was provided by the Center for Bioenergy Innovation (CBI) led by Oak Ridge National Laboratory. CBI is funded as a U.S. Department of Energy Bioenergy Research Centers supported by the Office of Biological and Environmental Research in the DOE Office of Science under FWP ERKP886. Oak Ridge National Laboratory is managed by UT-Battelle, LLC for the U.S. Department of Energy under contract no. DE-AC05-00OR22725. This work was also supported by the Genomics-Enabled Plant Biology for Determination of Gene Function program by the Office of Biological and Environmental Research in the DOE Office of Science (award DE-SC0020164). Research in the laboratory of GT is funded by the John B Orr Endowed chair in Environmental Plant Sciences. MCK acknowledges the Department of Plant Sciences, UC Davis, for the award of a GSR scholarship funded by endowments, particularly the James Monroe McDonald Endowment, administered by UCANR. MCK also gratefully acknowledges the UCD Jastro-Shields Research Award for their support.

